# 5’ tRNA halves are highly expressed in the primate hippocampus and sequence-specifically regulate gene expression

**DOI:** 10.1101/774380

**Authors:** Julia Jehn, Jana Treml, Svenja Wulsch, Benjamin Ottum, Verena Erb, Charlotte Hewel, Roxana N. Kooijmans, Laura Wester, Isabel Fast, David Rosenkranz

## Abstract

Fragments of mature tRNAs have long been considered as mere degradation products without physiological function. However, recent reports show that tRNA fragments (tRFs) play prominent roles in diverse cellular processes across a wide spectrum of species. Contrasting the situation in other small RNA pathways the mechanisms behind these effects appear more diverse, more complex and are generally less well understood. In addition, surprisingly little is known about the expression profiles of tRFs across different tissues and species. Here, we provide an initial overview of tRF expression in different species and tissues, revealing very high tRF-levels particularly in the primate hippocampus. We further modulated the regulation capacity of selected tRFs in human cells by transfecting synthetic tRF mimics (“overexpression”) or antisense-RNAs (“inhibition”) and identified differentially expressed transcripts based on RNAseq. We then used a novel k-mer mapping approach to dissect the underlying targeting rules, demonstrating that 5’ tRNA halves (5’ tRHs) silence genes in a sequence-specific manner, while the most efficient target sites align to the mid-region of the 5’ tRH and are located within the CDS or 3’ UTR of the target. This amends previous observations that tRFs guide Argonaut proteins to silence their targets via a miRNA-like 5’ seed match and suggests a yet unknown mechanism of regulation. Finally, our data suggests that some 5’ tRHs are also able to sequence-specifically stabilize mRNAs as upregulated mRNAs are also significantly enriched for 5’ tRH target sites.

## Introduction

tRNAs are well known for their conserved role in protein biosynthesis. However, mature tRNAs and their precursors give also rise to a class of small non-coding RNAs, the tRNA fragments (tRFs). While tRF1s (15-22 nt) are generated by clipping off the trailer sequence from the tRNA precursor molecule, 5’ tRFs and 3’ tRFs (18-35 nt) stem from the respective end of mature tRNAs (70-90 nt). Primarily considered as degradation products, tRFs are now recognized as additional players in small RNA-mediated gene regulation that act in a variety of cellular processes across species from all domains of life.

tRFs were found to play a role in fundamental physiological processes such as proliferation (Lee et al. 2009) and protein translation control (Gebetsberger et al. 2017; Guzzi et al. 2018; Ivanov et al. 2011; Keam et al. 2017). Moreover, tRFs are implicated in defense mechanisms of *Escherichia coli* against bacteriophages (Levitz et al. 1990) and human cells against trypanosoma or viruses (Deng et al. 2015; Garcia-Silva et al. 2014). Additionally, tRFs were found to prime the reverse transcription or inhibit the promotion of retroviruses and retrotransposons (Martinez et al. 2017; Ruggero et al. 2014; Schorn et al. 2017; Yeung et al. 2009). Furthermore, tRFs are associated with several diseases such as cancer (Goodarzi et al. 2015; Huang et al. 2017; Lee et al. 2009) and amyotrophic lateral sclerosis (Greenway et al. 2006; Ivanov et al. 2014) and were lately revealed to act as transgenerational transmitters that induce metabolic disorders and addictive behavior in mice (Chen et al. 2016; Sarker et al. 2019; Sharma et al. 2016; Short et al. 2017). While some of the described effects seem to base on similar mechanisms, tRFs from the same class were shown to trigger completely converse functions in other cases (Jehn and Rosenkranz 2019), suggesting that the regulatory potential of tRFs is more complex than observed for the well-studied major small RNA classes.

So far, two superordinate concepts that aim explain how tRFs regulate gene expression have been identified. While some studies show that tRFs globally repress translation by inhibiting the assembly of the translation initiation machinery (Gebetsberger et al. 2017; Guzzi et al. 2018; Ivanov et al. 2011; Keam et al. 2017), others report a sequence-specific gene regulation (Deng et al. 2015; Haussecker et al. 2010; Kuscu et al. 2018; Luo et al. 2018). As tRFs were found to associate with Argonaut proteins, a miRNA-like gene regulation mechanism seems apparent (Kumar et al. 2014; Kuscu et al. 2018; Maute et al. 2013). Indeed, studies could show that tRF-mediated transcript silencing is dependent on a 5’ seed, which is complementary to target sites within the 3’ UTR (Haussecker et al. 2010; Kuscu et al. 2018). Contrasting this, other studies observed sequence-specific gene silencing effects, where complementarity of the 5’ seed was neglectable in favor for other tRF portions (Deng et al. 2015; Luo et al. 2018). Since Argonaut structure coerces the 5' end of small RNAs for target recognition (Boland et al. 2011), the results of these studies suggest that tRFs interact with other effector proteins or act protein-independent to regulate genes. Indications from recent studies where tRFs were shown to interact with different proteins depending on the differentiation state of the cell or the modification status of the tRF further erode the AGO-centric view (Guzzi et al. 2018; Krishna et al. 2019) illustrating that we still lack a sufficiently deep understanding of how tRFs regulate gene expression mechanistically.

Surprisingly, although it has been noted that tRFs are much more abundant in tissues than in cultured cells (Torres et al. 2019), their expression profile across tissues and species has not yet been investigated. We therefore initially analyzed several available and own small RNA sequencing datasets to provide a first overview on tRF expression profiles. Subsequently, we tested if 5’ tRHs regulate genes by modulating the level of 5’ tRHs-Glu-CTC and 5’ tRH-Gly-GCC in cultured cells by transfecting either synthetic RNA mimics or inhibiting antisense RNAs. By RNA sequencing, *in silico* target predictions and a novel k-mer mapping-based approach, we then dissected the targeting rules of the respective 5’ tRHs. Finally, applying the k-mer analysis on similar RNA sequencing datasets of other species, we examined whether the identified targeting patterns of individual tRFs are conserved across species.

## Results and Discussion

### 5’ tRHs are highly abundant in the hippocampus of primates

To investigate the expression levels of tRFs across different tissues, we annotated available human small RNA sequencing datasets complemented by own small RNA sequencing data and found 5’ tRHs to be predominantly expressed in the hippocampus (figure 1A). Here, 30% of all mapped reads were classified as tRNA-fragments with 82% of these tRF-annotated sequences being 5’ tRHs (figure 1B). In comparison, the overall tRF level was only 13% in both the frontal cortex and the cerebellum (figure 1 B and C). Amongst the predominant tRNA-derived sequences in the hippocampus were the 5’ tRNA-halves 5’ tRH-Glu-CTC (26%) and 5’ tRH-Gly-GCC (10%). Notably, these 5’ tRHs were also amongst the most abundant 5’ tRHs in the analyzed small RNA sequencing libraries of other tissues (figure 1A). Both, 5’ tRH-Glu-CTC and 5’ tRH-Gly-GCC have been previously shown to play major roles in various cellular functions. They were amongst the sperm small RNAs that are involved in the epigenetic inheritance of paternal diet-induced metabolic disorders and addictive-like behavior in mice (Chen et al. 2016; Sarker et al. 2019; Sharma et al. 2016; Zhang, Y. et al. 2018). Additionally, they were found amongst a group of 5’ tRHs to be upregulated in cells upon infection by the Respiratory Syncytial Virus (Wang et al. 2013). 5’ tRH-Gly-GCC was furthermore found to be part of a specific subset of 5’ tRHs that is dynamically expressed during stem cell differentiation (Krishna et al. 2019), while 5’ tRH-Glu-CTC was found to be highly expressed in human monocytes, where it triggers the transcriptional suppression of the surface glycoprotein CD1 (Zhang et al. 2016).

**Figure 1:**
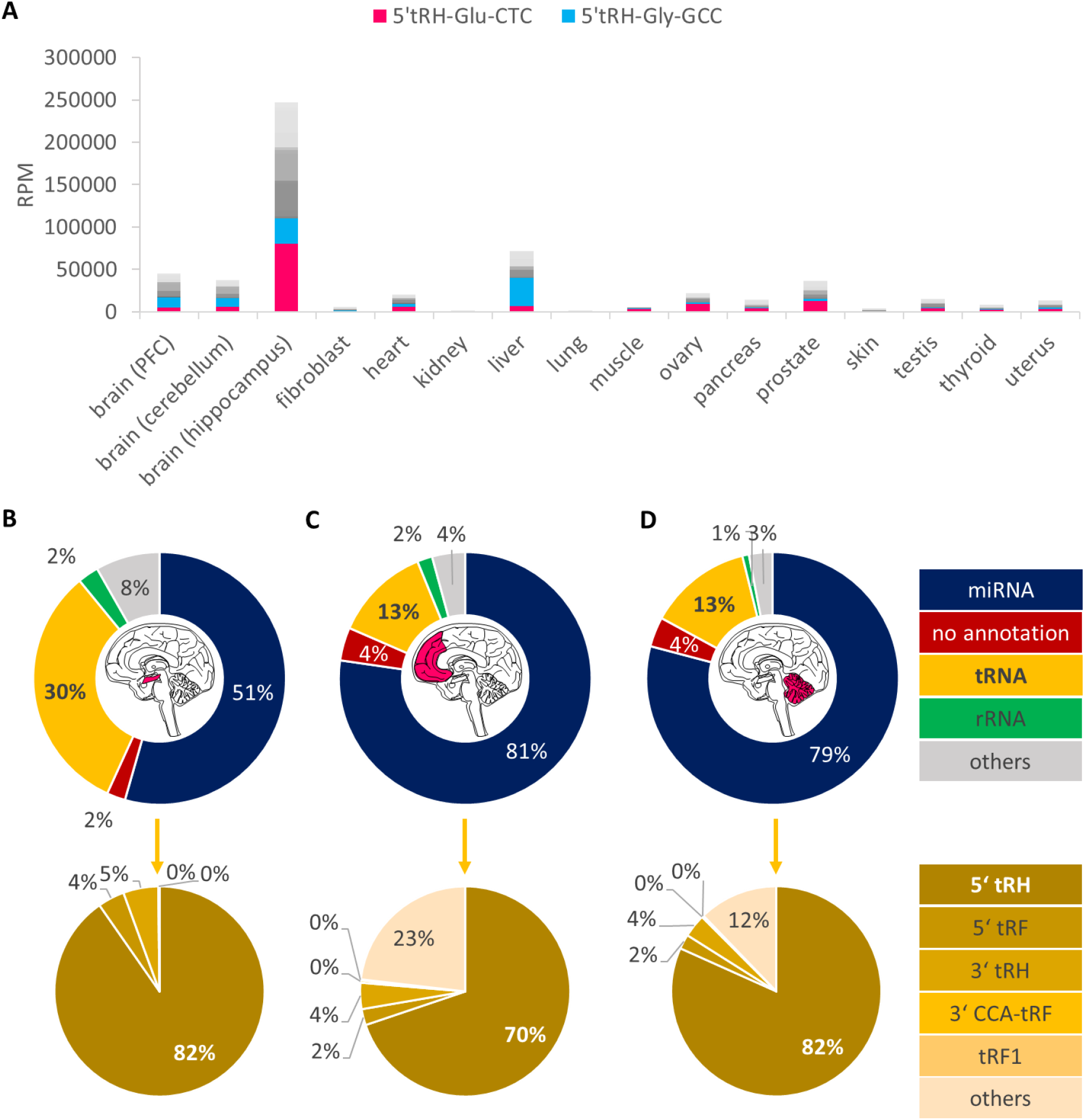
**(A)** Reads per million (RPM) of small RNAs annotated as 5’ tRHs in small RNA sequencing datasets of human tissues. The mean value of multiple datasets is shown. **(B-D)** Small RNA annotation of mapped reads from small RNA libraries of the human hippocampus, frontal cortex and cerebellum.

In order to investigate whether high 5’ tRH levels in the hippocampus are a common feature in mammals, we generated small RNA sequencing libraries of chimpanzee and macaque hippocampus samples, and additionally analyzed publicly available small RNA sequencing datasets from pig, rat and mouse hippocampus. Interestingly, we found even higher proportions of reads being annotated as tRFs in the hippocampal libraries of the two primates (42% in the chimpanzee / 51% in the macaque; figure 2) compared to the human hippocampus libraries (30%). Again, 5’ tRHs made up the majority of the tRF-annotated reads (82% / 75%) and 5’ tRH-Gly-GCC (15% / 33%) and 5’ tRH-Glu-CTC (14% / 11%) were amongst the most abundantly expressed tRFs. In contrast, even though 5’ tRHs are also the predominant tRF class in the analyzed sequencing libraries of the other mammals, only 10% of the pig and 3% of the rodent mapped reads were classified as tRFs (figure 2). This suggests that high levels of 5’ tRHs in the hippocampus are a primate-specific trait, which raises the question if a conserved subset of 5’ tRHs specifically fine-tunes hippocampal gene expression in primates.

**Figure 2:**
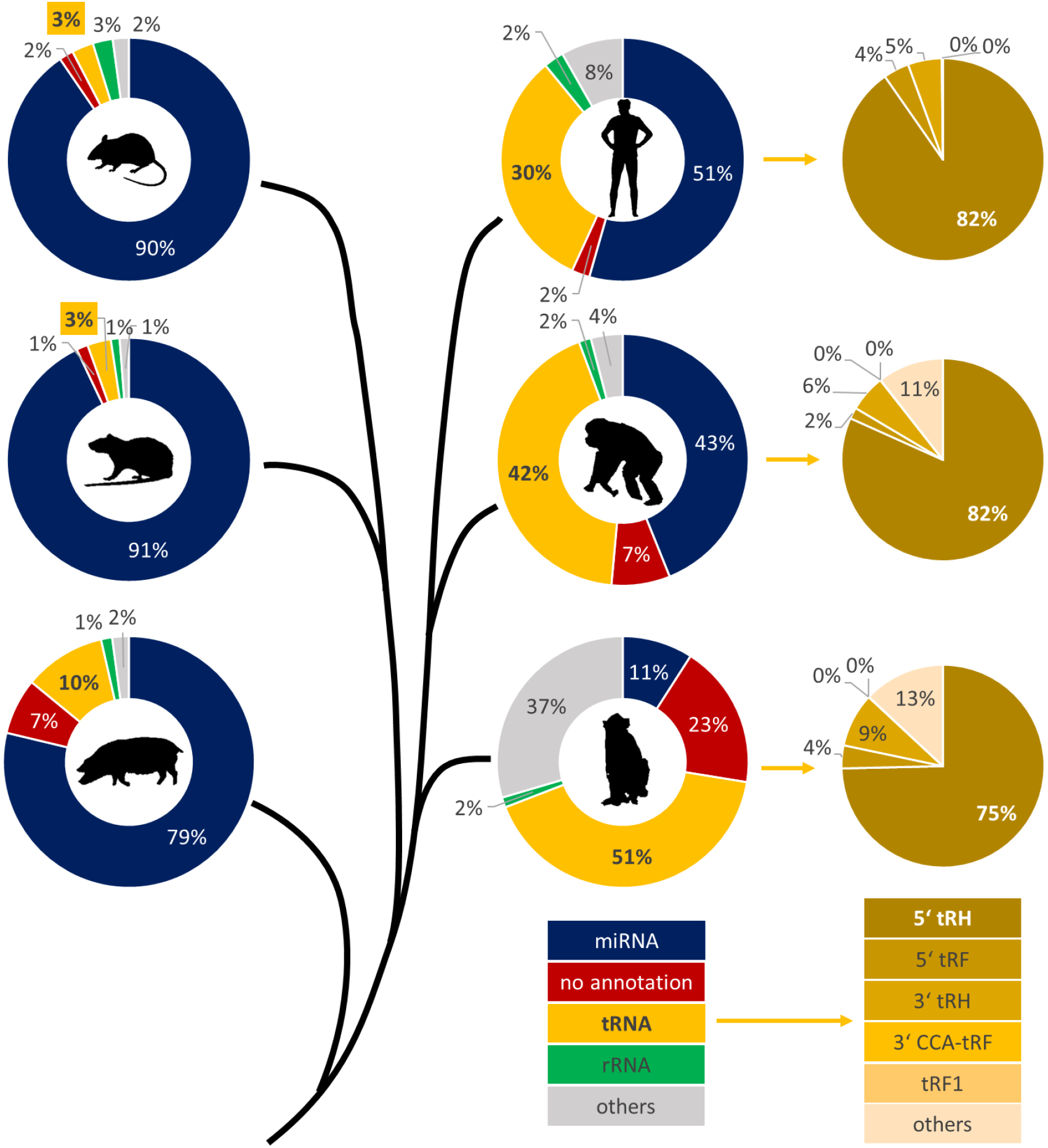
Phylogenetic tree and small RNA annotation of mapped reads from small RNA sequencing libraries of hippocampal samples from mouse, rat, pig, human, chimpanzee and macaque.

### miRNA- and piRNA-like targeting rules scarcely identify targets of the 5’ tRNA-halves Glu-CTC and Gly-GCC in HEK293T

In order to identify genes that are regulated by these 5’ tRHs, we overexpressed 5’ tRH-Glu-CTC and 5’ tRH-Gly-GCC in HEK293T cells by transfecting 50 nM synthetic 5’ tRH mimics. HEK293T cells were chosen as 5’ tRHs are barely expressed in this cell line: from the 9% of all mapped reads that were assigned as tRNA-fragments, only 1% are 5’ tRHs (Supplemental_Fig_S1A). As validated by quantitative RT-PCR (qPCR), transfection of the RNA mimics successfully increased the number of 5’ tRH copies in comparison to a non-target-siRNA transfection control (Supplemental_Fig_S2A).

As a first test, we quantified the expression change of twenty abundantly expressed transcripts, which were predicted to represent targets of the respective tRFs according to miRNA (miRanda) or piRNA targeting rules, via qPCR. For piRNA-like target prediction, we developed a software named piRanha that by default applies targeting rules empirically verified by Zhang et al. (Zhang, D. et al. 2018). All ten tested potential targets of 5’ tRH-Gly-GCC were lower expressed in the overexpression cells compared to the control cells. In contrast, neither the five miRanda- nor the five piRanha-predicted 5’ tRH-Glu-CTC-target transcripts were differentially expressed upon overexpression of the 5’ tRH-Glu-CTC (figure 3A), suggesting that target rules might differ between different tRFs.

**Figure 3:**
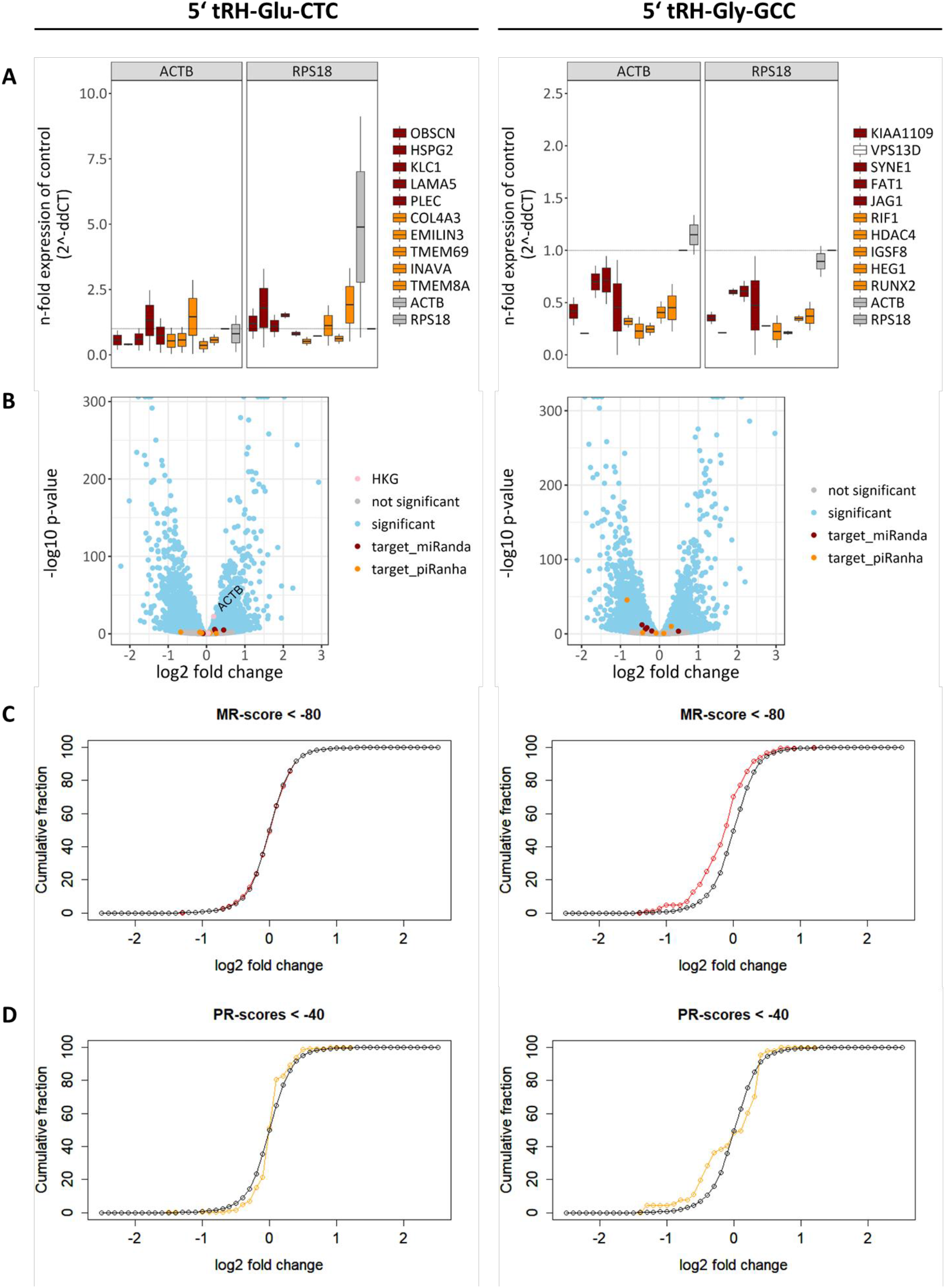
**(A)** n-fold expression of potential 5’ tRH targets in 5’ tRH overexpression HEK293T cells compared to control cells as quantified by qPCR. The selected transcripts were predicted by miRanda (red) and piRanha (orange) to be targets of the 5’ tRNA-halves Glu-CTC (left) or Gly-GCC (right). The housekeeping genes β-actin and RPS18 were used as normalizers. Note that either β-actin or RPS18 expression changes upon transfection of the 5’ tRNA-half Glu-CTC. **(B)** Volcano plot of differential expression analysis for protein-coding genes of 5’ tRH overexpression and control HEK293T cells (blue, adjusted p-value < 0.01). The 10 previously tested putative target genes are highlighted in red (miRanda prediction) and orange (piRanda prediction). For the 5’ tRH-Glu-CTC analysis (left), the housekeeping genes β-actin and RPS18 are additionally highlighted in pink, showing that β-actin (ACTB) gets upregulated upon 5’ tRH-Glu-CTC overexpression, while the RPS18 expression is not affected. **(C/D)** Cumulative plots for log2-fold-change values of genes that were identified as potential targets of the 5’ tRHs Glu-CTC or Gly-GCC by miRanda (red) or piRanha (orange) and of all genes (black). Only in 5’ tRH-Gly-GCC overexpression HEK293T cells miRanda-predicted genes are significantly more repressed.

To get a global overview of 5’ tRH-regulated genes, we then sequenced the transcriptome of HEK293T 5’ tRH-overexpression and control cells and performed differential expression analysis. Upon overexpression of the respective 5’ tRH, more genes were significantly downregulated than upregulated (adjusted p-value < 0.01). However, most of the potential 5’ tRH targets previously quantified by qPCR were not significantly differentially expressed (figure 3B).

When analyzing the cumulative distribution of all miRanda and piRanha predicted genes, only miRanda-targets of 5’ tRH-Gly-GCC were enriched for downregulated genes over the general distribution (figure 3C and D). We therefore conclude that for 5’ tRNA-halves, the targeting rules of piRNAs are not applicable, while the targeting rules of miRNAs cannot satisfyingly predict 5’ tRH targets neither.

### Non-miRNA-like targeting rules for 5’ tRHs

To unravel the targeting rules of 5’ tRHs, we developed a k-mer mapping approach, where k-mers of the 5’ tRH sequences with all possible lengths (k ≥ 5) and start positions within the tRH are mapped separately to the three major regions of each human transcript (5’ UTR, CDS and 3’ UTR). By calculating the fraction of “targeted” (k-mer alignment) or “not targeted” (no k-mer alignment) genes per k-mer that get significantly downregulated (adjusted p-value < 0.01) we identify the portion within the tRH that is most likely to be important for target recognition and thus silencing. Figure 4 visualizes the underlying principles of the analysis.

**Figure 4:**
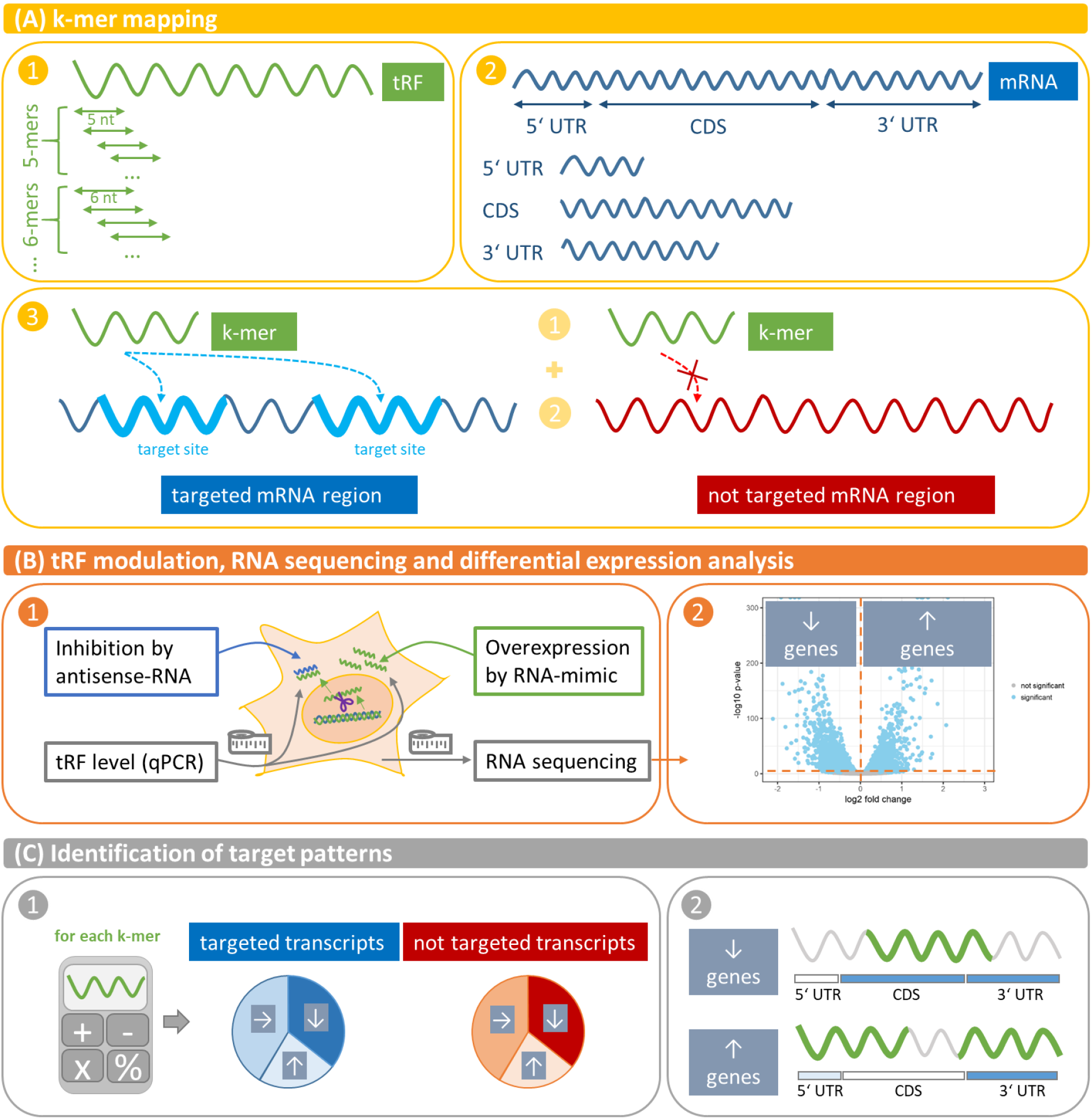
Graphical overview of identification of tRF targeting rules via a k-mer mapping approach.

Our analysis revealed that downregulation of transcripts with k-mer alignment (“targeted”) is significantly enriched over the number of downregulated transcripts without k-mer alignment (“not targeted”) particularly when k-mers of 5-10 nt length with start positions in the middle of the 5’ tRH align to the 3’ UTR or the CDS of the target (figure 5). Thus, unlike miRNAs that bind to the 3’ UTR of their targets via a 7 nt long seed at the 5’ end, 5’ tRNA-halves such as 5’ tRH-Glu-CTC and 5’ tRH-Gly-GCC downregulate transcripts mainly by binding with a 5-10 nt long stretch of their middle region to the 3’ UTR or the CDS of the target transcripts.

**Figure 5:**
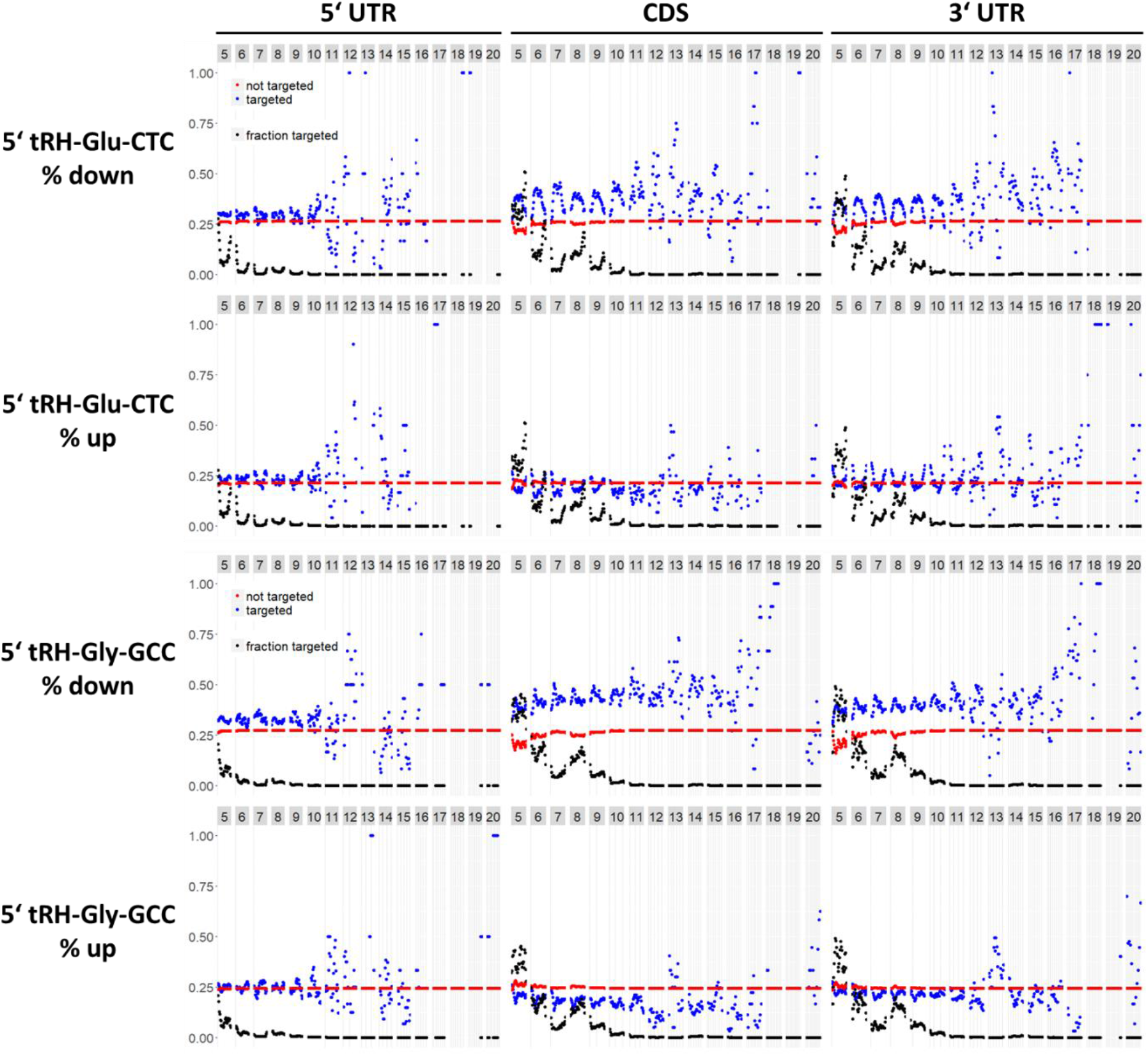
k-mer analysis to elucidate the targeting rules of 5’ tRH-mediated transcript regulation in HEK293T cells. The plots shows for each k-mer the percentage of transcripts with (blue) and without k-mer alignment (red) that are down/-upregulated. The fraction of genes targeted by the respective k-mer is displayed in black.

The resulting target pattern additionally suggests that 5’ tRH-Gly-GCC, but not 5’ tRH-Glu-CTC, is able to downregulate targets by binding with its miRNA-seed like 5’end to the transcript. This is in line with, the different target regulation behavior of 5’ tRH-Glu-CTC and 5’ tRH-Gly-GCC as revealed by qPCR quantification of target transcripts predicted with miRanda.

As a previous study found that 5’ tRH-Glu-CTC interacts with PIWIL4 to transcriptionally silence the surface glycoprotein CD1 in human monocytes (Zhang et al. 2016), we wanted to check whether we find indications for transcriptional silencing by 5’ tRH-Glu-CTC and 5’ tRH Gly-GCC in our datasets. We therefore restricted our k-mer analysis to alignments that span target gene exon-junctions. If the corresponding 5’ tRHs suppresses their target genes at the transcriptional level, we would expect a decreasing fraction of transcripts that have an exon-junction spanning alignment and that are downregulated. In contrast, the fraction of transcripts that have a non-exon-junction spanning alignment and that are downregulated should not change. As we did not observe such an effect, we conclude that the studied 5’ tRHs silence their targets at the post-transcriptional level in HEK293T cells (Supplemental_Fig_S3).

Interestingly, mRNA quantification by qPCR and RNA sequencing revealed that the housekeeping gene *ACTB* gets upregulated upon 5’ tRH-Glu-CTC transfection (figure 3A and B). Unexpectedly, *ACTB* has several potential binding sites for 5’ tRH-Glu-CTC. Hence, we were curious to check whether upregulated genes might be enriched for 5’ tRHs target sites as well, which would indicate that 5’ tRH targeting might also have a protective effect. Indeed, our analysis revealed that transcripts that have target sites in their 3’ UTR for the 5’ and 3’ ends of 5’ tRH-Glu-CTC are more likely to be upregulated than those that do not have a target site, while a corresponding upregulating effect was not detectable for 5’ tRH-Gly-GCC (figure 5).

Concluding from the above outlined target pattern analyses we assume that tRNA-fragments, even from the same tRF-series, may recognize their targets via different parts of the tRF instead of a dominating 5’ seed match (figure 5). This demonstrates that a Argonaut-dependent regulation mechanism as proposed for tRFs previously (Haussecker et al. 2010; Kuscu et al. 2018) cannot fully explain the observed changes in gene expression. However, our results are in line with other studies identifying miRNA-untypical targeting (Deng et al. 2015; Luo et al. 2018). The mechanism seems to act at the post-transcriptional level, as transcripts with exon-junction spanning k-mer alignments were similarly often downregulated compared to transcripts with intra-exonic k-mer alignment (Supplemental_Fig_S3). Our analysis additionally suggests that some tRFs (e.g. 5’ tRH-Glu-CTC) trigger not only the downregulation of target genes, but may also stabilize target transcripts (figure 5). This is a rather unexpected finding, since a stabilizing effect of small RNAs is rarely seen in eukaryotes, but rather a trait of prokaryotic small RNA pathways (Fröhlich and Vogel 2009). However, a recent study showed that sequence-specific binding of a 3’ tRF to the mRNAs of ribosomal proteins enhances the translation of the target transcript (Kim et al. 2017). It was suggested that structural changes induced by this interaction allow for higher transcription rates. Similar alterations in the secondary structure could also lead to a stabilization of the transcript.

### Identification of genuine 5’ tRH targets by antisense-inhibition of 5’ tRH-Glu-CTC suggests a role of 5’ tRHs in neurogenesis

Noteworthiliy, drawing conclusions regarding tRF targets only from overexpression experiments has several shortcomings. First, our synthetic RNA mimic does not carry post-transcriptional modifications as endogenous tRFs do. Thus, we might observe an artificial regulation behavior that does not reflect the physiological situation. Second, genes that were found to be differentially expressed upon tRF overexpression in cells that typically express these tRFs at very low levels must not necessarily be genuine targets of the modulated tRF under natural conditions. To circumvent these distorting effects and gain support for our conclusion, we additionally performed an inverse experiment where we inhibited the regulation capacity of 5’ tRH-Glu-CTC by transfecting an antisense RNA.

We chose to perform the experiment with HepG2 cells, in order to exclude cell line specific regulation effects. HepG2 cells have a similar overall tRF level compared to HEK293T cells (6% of the total reads compared to 9% in HEK293T cells; Supplemental_Fig_S1B), but express a higher proportion of 5’ tRHs according to small RNA sequencing data (17% of the reads assigned to tRNAs compared to 1% in HEK293T cells; Supplemental_Fig_S1B). Like in HEK293T cells, 5’ tRH-Glu-CTC and 5’ tRH-Gly-GCC are the most abundant 5’ tRHs (6% and 3% of the reads assigned to tRNAs) in the analyzed library. As quantified by qPCR, transfection of the antisense-RNA decreased the number of 5’ tRH-Glu-CTC copies in HepG2 cells by about 70% (Supplemental_Fig_S2B). Even though the level of 5’ tRH-Glu-CTC was decreased, only a small amount of genes was significantly differentially expressed compared to the control state (818 downregulated and 717 upregulated genes, adjusted p-value < 0.01; figure 6A). While the majority of significantly differentially expressed genes of the antisense-inhibited HepG2 was likewise regulated as in the overexpression HEK293T cells (figure 6B grey areas), only few genes were inversely regulated (figure 6B rosy and lutescent area) and we assume these genes to represent the genuine targets of 5’ tRH-Glu-CTC. In the following we refer to the 34 genes that might get downregulated by 5’ tRH-Glu-CTC as “perish targets” (figure 6B rosy area) and name the subset of 50 genes that get upregulated possibly due to stabilizing effects exerted by 5’ tRH-Glu-CTC “shelter targets” (figure 6B lutescent area).

**Figure 6:**
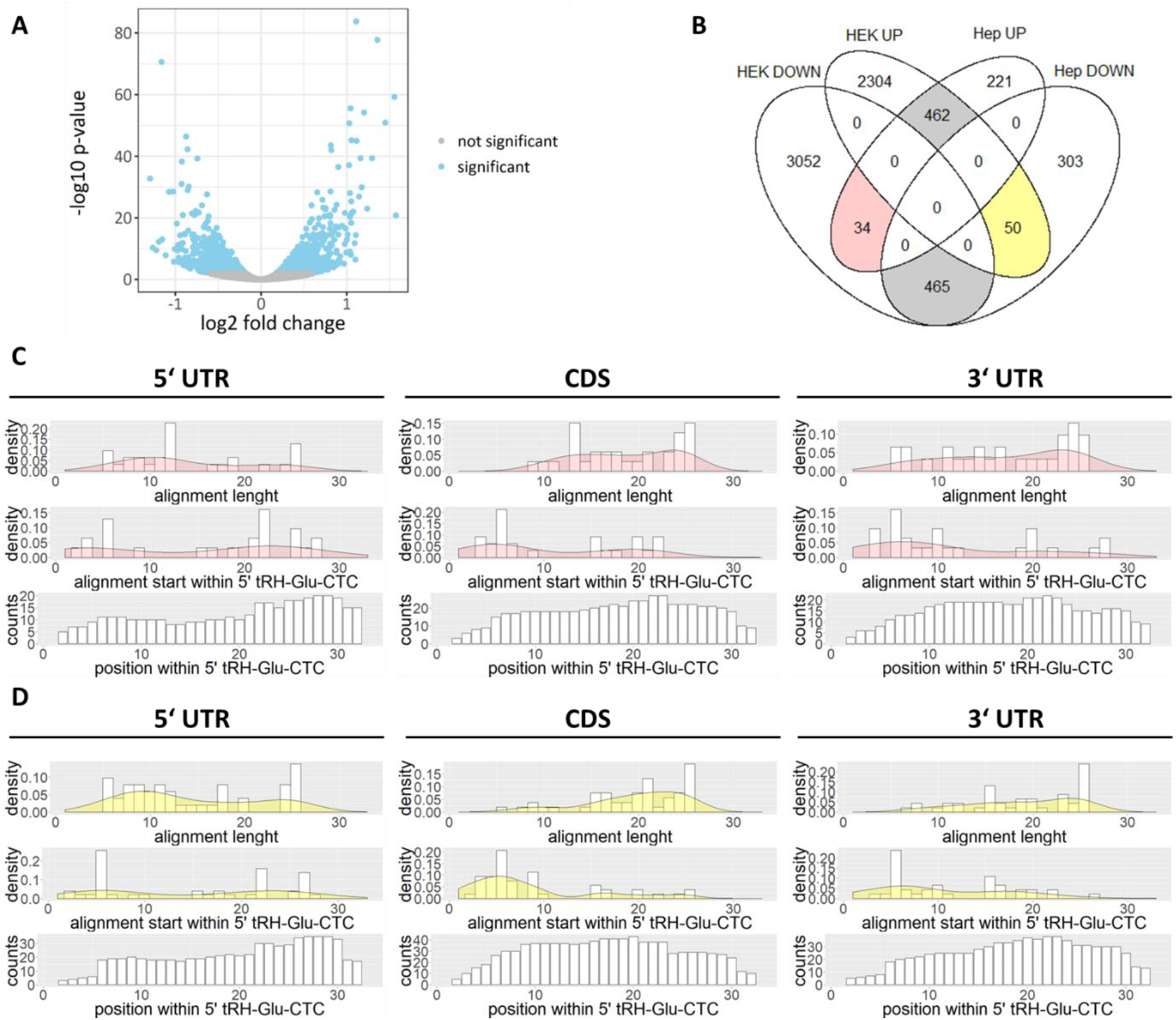
**(A)** Volcano plot of differential expression analysis for protein-coding genes of 5’ tRH antisense-inhibition and control HepG2 cells (blue, adjusted p-value < 0.01). **(B)** Venn diagram of significantly differentially expressed genes. Genes that are likewise regulated in HEK as in HepG2 cells are highlighted in grey. Genes that are inversely regulated in HEK overexpression and HepG2 inhibition cells are highlighted rosy (“potential perish targets”) and lutescent (“potential shelter targets”). **(C/D)** Analysis of thermodynamically favored alignments for the major transcript regions of “potential perish targets” or “potential shelter targets” with 5’ tRH-Glu-CTC.

In order to characterize 5’ tRH-targeting further, we subsetted the three major transcript regions (5’ UTR, CDS, 3’ UTR) of the potential perish and shelter targets and computed the thermodynamics for RNA-duplex formation with the 5’ tRH-Glu-CTC. Considering the free energy needed to open intrinsic secondary structures, the free energy of the interaction and the so-called “dangling end” energies that non-interacting sequence stretches are causing, we identified the energetic optimal region of interaction. For the CDS and the 3’ UTR of both the potential perish targets and the potential shelter targets, these optimal binding sites were enriched for k-mer alignments with a length of around 20 bp (figure 6C and D) and alignments preferentially started around the sixth position from the 5’ end of the tRH. Thus, most thermodynamically favored interactions involved big stretches of the middle part of 5’ tRH-Glu-CTC. This is in line with the target pattern identified by the k-mer mapping approach in case of the potential perish targets, but not with the target pattern for the potential shelter targets. As stabilizing effects probably involve other protein interactors than silencing effects, this objection may be a result of the stabilizing interactor exposing only the ends of the tRH for target recognition.

As we found 5’ tRHs like 5’ tRH-Glu-CTC to be extremely abundant in small RNA libraries of primate hippocampal tissues and previous studies suggested a role in targeting genes involved in neural processes (Krishna et al. 2019; Sarker et al. 2019), we were interested whether the identified potential targets of 5’ tRH-Glu-CTC are implicated in neuronal processes. Indeed, we found 20% of the potential shelter targets (transcripts might be stabilized), but only 5% of the potential perish targets (transcripts might be degraded) to have an assigned neuronal function. In comparison only 7% and 8% of all expressed genes in HEK293T and HepG2 cells were assigned the gene ontology term “neurogenesis” (GO:0022008). Hence, there is a statistic correlation between genes that are potential targets of 5’ tRH-Glu-CTC and neurogenesis (α<0.001 for shelter targets and α<0.01 for perish targets; chi-squared test). While genes like *MDK*, *VEGFA* and *EVL* play a role in the regulation of axon outgrowth (Drees and Gertler 2008; He et al. 2015; Kurosawa et al. 2001), genes like *NOTCH1* and *NP1L1* are involved in neuronal differentiation (Patten et al. 2006; Qiao et al. 2018). Interestingly, decapitation studies with the planarian *Dugesia japonica* showed that 5’ tRH-Gly-GCC is not only upregulated in regenerating animals, but is furthermore required for proper head regeneration (Z Cao, D Rosenkranz, S Wu, H Liu, Q Pang, B Liu, B Zhao; manuscript in preparation). Taking into consideration that planarians share many CNS genes with humans (Mineta et al. 2003), and that the hippocampus is one of the few brain regions known to have high rates of adult neurogenesis (Boldrini et al. 2018; Eriksson et al. 1998), it is tempting to speculate that 5’ tRHs may have critical functions in human neurogenesis as well.

### Targeting pattern of 5’ tRHs/tRFs are unique for each fragment, but seem to be conserved across species

In order to gain further support for our assumptions, we checked whether the identified 5’ tRH target patterns are conserved among different species. We therefore analyzed published RNA sequencing data from Drosophila S2 cells that were transfected with a 20 nt long 5’ tRF-Glu-CTC, which exhibits high sequence homology with the human 5’ tRNA-half (Supplemental_Fig_S4). As is the case for the human 5’ tRNA-half, the fly 5’ tRNA-fragment of the tRNA-Glu-CTC showed the strongest downregulating effect on targets when its ~7 nt long middle part binds to the 3’ UTR or CDS of the target transcript (figure 7). However, unlike the human 5’ tRNA-half, the both ends of the fly 5’ tRF did not seem to contribute to target upregulation (data not shown), which is possibly due to its shorter sequence which lacks the corresponding nucleotides. Given the similar target regulation pattern, we suggest that 5’ tRNA-fragments can regulate their targets via conserved mechanisms across different species, while the sequence stretch being most important for target regulation can vary for fragments from different parental tRNAs.

**Figure 7:**
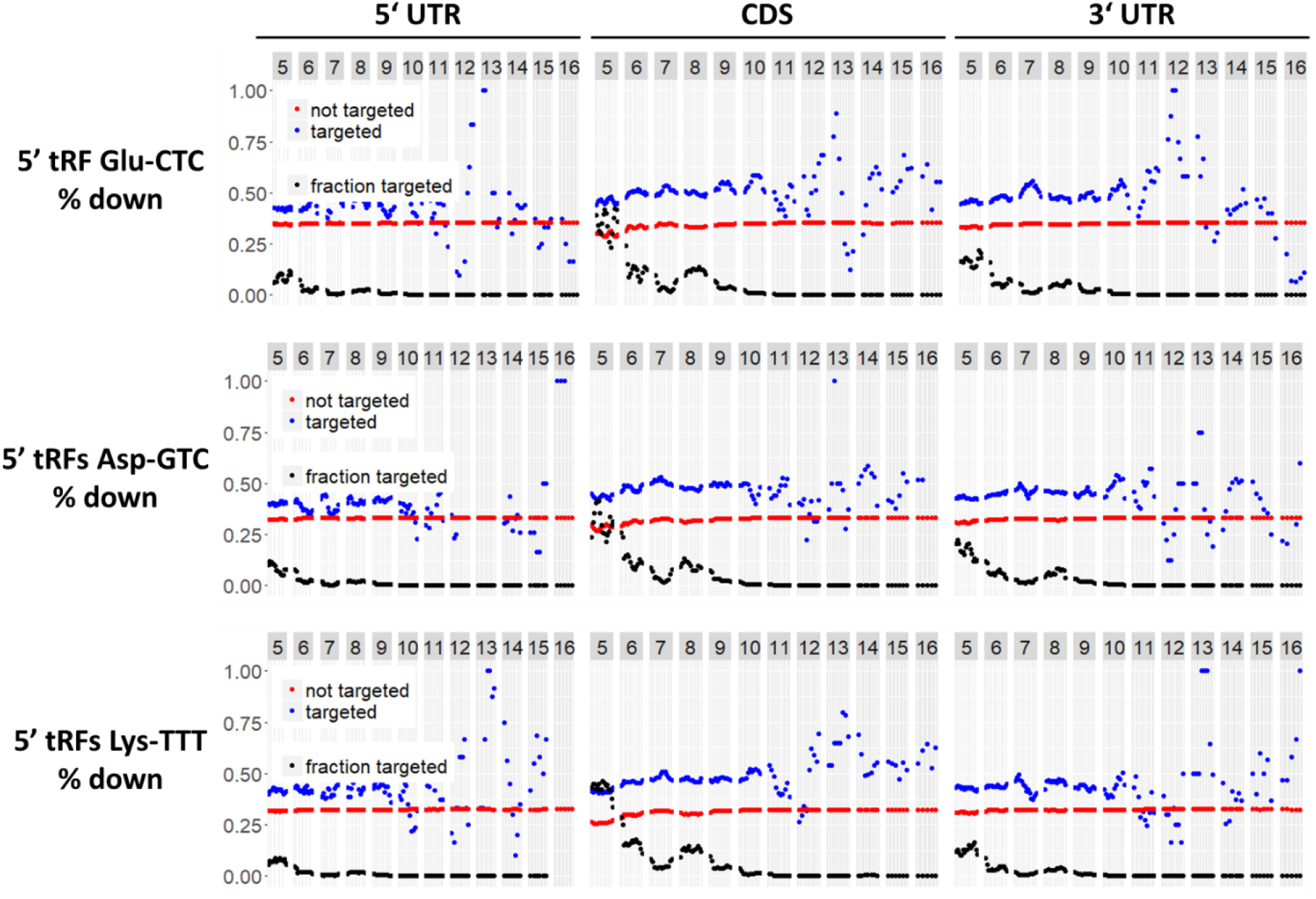
Target pattern analysis of published RNA sequencing data of fly S2 cells, where the 20 nt long 5’ tRF Glu-CTC, 5’ tRFs Asp-GTC or Lys-TTT was overexpressed by tRF mimic transfection (Luo et al. 2018). Plotted is the percentage of genes with (blue) or without (red) k-mer alignment that get downregulated.

To confirm this assumption, we analyzed additional published RNA sequencing datasets from fly S2 cells that were transfected with the 20 nt long 5’ tRNA-fragments Asp-GTC and Lys-TTT (figure 7). These analyses support our hypothesis, that 5’ tRNA-fragments from different parental tRNAs act via different regions to regulate their targets on a sequence-complementary basis. What they have in common, is that 3’ UTR and CDS targeting leads to the biggest regulatory effect.

### Concluding remarks

Despite the rise of NGS techniques, exploring the mechanisms by which small RNAs target genes remains challenging. Even for miRNAs, where gene regulation mechanisms are well studied, correct target prediction is difficult since the *in vivo* accessibility of potential target sites is difficult to assess. RNA binding proteins may not only occupy the putative binding site, but may also change the secondary structure of the target when binding elsewhere in the transcript. Predicting and identifying targets of rather unexplored small RNAs like tRFs is the more complicated, as it is unknown if and to which extend mismatches, wobble base pairs and bulges are tolerated. Aggravatingly is that target interactions can be surprisingly variable (Backofen and Hess 2010).

Using a k-mer mapping approach, we sought to identify the target patterns of 5’ tRHs. Our analysis suggested that 5’ tRHs silence genes, which have complementary binding sites for long stretches of the tRNA half that not necessarily need to include the miRNA-typical 5’ region. This finding strongly suggests that Argonaut proteins are not necessarily indispensable as effector proteins for tRF-dependent gene regulation as it had been suggested by other studies (Haussecker et al. 2010; Kuscu et al. 2018). Whether 5’ tRHs independently regulate targets, or do so in association with other effector proteins than Argonauts remains to be investigated. As it was shown that specific subsets of tRFs bind to certain proteins (Goodarzi et al. 2015; Ivanov et al. 2011; Krishna et al. 2019; Saikia et al. 2014), it is likely that 5’ tRHs might interact with different proteins to regulate distinct targets. The tRNA 3’ processing endoribonuclease RNase Z might be such an effector protein, as it was shown to cleave transcripts, which form RNA hybrids with 5’ tRHs that have similar secondary structures like pre-tRNAs (Elbarbary et al. 2009). Identifying more proteins involved in tRF-mediated gene regulation and elucidating the underlying mechanism will greatly enhance our understanding of gene regulation.

Analyzing the expression profile of tRFs across tissues and species, we found 5’ tRHs to be particularly high expressed in the hippocampus of primates, while their expression was rather low in the hippocampus of the pig, the rat and the mouse. In the hippocampus of all three tested primate species (human, chimpanzee and macaque), 5’ tRH-Glu-CTC and 5’ tRH-Gly-GCC were amongst the most abundant 5’ tRHs suggesting that they have a conserved role in the primate hippocampus. As we find transcripts that are presumably stabilized by 5’ tRH-Glu-CTC targeting to be substantially enriched for a function in neuronal processes such as axon outgrowth and neuronal differentiation, while transcripts that are presumably degraded upon 5’ tRH-Glu-CTC targeting are rather depleted of a function in neurogenesis, we hypothesize that 5’ tRHs play a role in fine-tuning primate neurogenesis. Alternatively, since retrotransposons are highly active in hippocampal neurons (Upton et al. 2015) and as 5’ tRH-Gly-GCC was shown to downregulate transcripts that are driven by the LTR-retrotransposon MERVL in the developing mouse embryo (Sharma et al. 2016), it is also possible that hippocampal 5’ tRH expression serves the purpose of regulating transposition-related events.

## Materials and Methods

### Small RNA sequencing, data processing and annotation

Small RNA sequencing data of various human tissues was downloaded from the SRA Database (for accession numbers see Supplemental Table Supplemental_Table_S1) and quality checked by FastQC (version 0.11.2). Adapter and quality trimming was performed using BBDuk (version 36.77; ktrim=r overwrite=true k=20 mink=9 ziplevel=2 hdist=1 qtrim=rl trimq=10 minlen=15 maxlen=34; for Encode data additionally: forcetrimleft=6) before the reads were FastQC checked again and mapped to the human genome (version GCA_000001405.27_GRCh38.p12) using Bowtie 2 (version 2.3.0). Based on these map-files small RNA annotation was performed with unitas (version 1.7.3; Gebert et al. 2017). Using the custom Perl script annotationtable2RPM.pl the RPM values were calculated for the respective tRF species.

Total RNA from adult normal male human hippocampal tissue was obtained from AMS Biotechnology (Cat. Nr: R1234052-10). A small RNA library was prepared as described in Gebert et al. 2015. In brief, small RNAs (15-40 nt) were extracted from a denaturing polyacrylamide gel. Subsequently, a 3’ adapter (5’-rAppCTGTAGGCACCATCAATddC-3’) and a 5’ adapter (5’-GACUGGAGCACGAGGACACUGACAUGGACUGAAGGAGUAGAAA-3’) were ligated to the small RNAs. Following cDNA synthesis with the RT-Primer 5’-ATTGATGGTGCCTACAG-3’, the cDNA transcripts were PCR amplified using the forward primer 5’-ACATGGACTGAAGGAGTAGA-3’ and the index-containing reverse primer 5’-ggctcATTGATGGTGCCTACAG-3’. The generated library was high throughput sequenced in parallel with six other indexed samples by GENterprise (Mainz) on an Illumina HiSeq 2000 system. After converting the fastq file to fasta formate using the NGS toolbox Perl script *fastq2fasta*, the 5’ adapter sequence was clipped off the 120 nt long reads using the NGS toolbox Perl script *clip* (−m 5.AGTAGAAA). As reads may contain the reverse complementary sequence, the reverse complementary variant of the 5’ adapter sequences were also clipped off (−m TTTCTACT.3) and the remaining sequences were transcribed to the original sequence direction using the NGS toolbox Perl script *rev-comp*. Both outputs were then concatenated using the NGS toolbox Perl script *concatenate*. To extract only the reads with the right index and to remove the 3’ adapter sequence the NGS toolbox Perl script *clip* (−m CTGTA.GAGCC.3) was applied. Subsequent analysis was performed as described below.

Total RNA from tissue samples of human adult brain regions (hippocampus, cortex, cerebellum) was obtained from the BioChain Institute (Newark, CA, USA). Total RNA of hippocampal tissue samples of a female and a male chimpanzee brain was obtained from the National Chimpanzee Brain Resource (www.chimpanzeebrain.org, USA). A macaque brain provided by the Primate Brain Bank (The Netherlands Institute for Neuroscience, Amsterdam, Netherlands) was dissected and hippocampal tissue was homogenized in TRIzol™ (Thermo Fisher). Total RNA was isolated according to the manufacturers protocol. The total RNA was sent to LC Sciences (Houston, TX, USA) for small RNA library preparation and small RNA sequencing. The 3’ adapter sequences were clipped off using the NGS toolbox (version 2.1; http://www.smallrnagroup.uni-mainz.de/) Perl script *clip* (-m TGGAATTC.3).

Small RNA sequencing data of hippocampal tissue from three male adult Wistar rats (Study: PRJEB24026, Run accessions: ERR2226477, ERR2226482 and ERR2226487) was downloaded from the European Nucleotide Archive. Subsequent analysis was performed as described below. Small RNA sequencing data of hippocampal tissue from the pig (SRR3105507 and SRR3105508) and the mouse (SRR5144167, SRR5144168 and SRR5144169) was downloaded via the SRA toolkit tool *fastq-dump* (version 2.8.2). The 3’ adapter sequences were clipped off using the NGS toolbox (version 2.1) Perl script *clip* (-m TGGAATTC.3). Subsequent analysis was performed as described below.

Total RNA of HEK293T cells was extracted with TRI Reagent™ (Thermo Fisher) 24 h after transfection with 10 nM Silencer™ Select Negative Control No. 1 siRNA (4390843; Thermo Fisher) using Lipofectamine™ RNAiMAX (Thermo Fisher). The RNA was sent to BGI (Hongkong) for small RNA library preparation and 50 bp single-end sequencing. The reads were delivered adapter-trimmed. Subsequent analysis was performed as described below.

Small RNA sequencing data of HepG2 cells (SRR6823987) was downloaded via the SRA toolkit tool *fastq-dump* (version 2.8.2). 3’ adapter sequences were clipped off using the NGS toolbox (version 2.1) Perl script *clip* (−m AGATCGGA.3). Subsequent analysis was performed as described below.

Adapter-trimmed data was first quality checked by FastQC (version 0.11.2), then converted to the fasta-format, length filtered (15-40 nt), collapsed to non-identical reads and depleted for low complexity reads using the NGS toolbox Perl scripts *length-filter*, *collapse* and *duster* (version 2.1; http://www.smallrnagroup.uni-mainz.de/). The remaining sequences were mapped to the respective genome (versions GCA_000001405.27_GRCh38.p12, GCA_000001515.5_Pan_tro_3.0, GCA_000772875.3_Mmul_8.0.1, GCA_000003025.6_Sscrofa11.1, GCA_000001895.4_Rnor_6.0 and GCA_000001635.8_GRCm38.p6) using the Perl script *sRNAmapper* (version 1.0.5; -a best) that employs SeqMap (Jiang and Wong 2008) as mapping tool. The map-files were used for small RNA annotation with unitas (version 1.6.1; Gebert et al. 2017).

### Transfection of tRF-mimics and tRF antisense 2’-OMe-RNAs

HEK293T (2.5E4 cells/well) and HepG2 (1E5 cells/well) cells were seeded in 24-well plates and cultured in 1x GlutaMAX™-I DMEM supplemented with 10% FBS (Thermo Fisher). The next day the cells were transfected with 50 nM tRF-mimics or tRF antisense 2’ OMe-RNAs (biomers) using Lipofectamine™ RNAiMAX (Thermo Fisher) according to the manufacturer’s protocol. Sequences of the transfected RNAs are available in Supplemental Table Supplemental_Table_S2. As control, cells were transfected with Silencer™ Select Negative Control No. 1 siRNA (Thermo Fisher). 48 hours after transfection, the RNA was isolated according to the TRI Reagent™ protocol (Thermo Fisher).

### RT-PCR quantification of tRFs

In order to measure the tRF level after transfection, the 15-40 nt small RNA fraction was eluted from a denaturing polyacrylamide gel (see Gebert et al. 2015) and polyadenylated using the A-Plus Poly(A) Polymerase Tailing Kit (Cellscript). After ethanol precipitation the poly-A tailed RNA was reversely transcribed with the SuperScript™ IV reverse transcriptase (Thermo Fisher) using the RT-primer 5’-CGAATTCTAGAGCTCGAGGCAGGCGACATGT25VN-3’. For qPCR 1 μL of cDNA was mixed 0.5 μL 10 μM sequence-specific forward primer, 0.5 μL 10 μM RT-primer-specific reverse primer, 3 μL water and 5 μL 2x QuantiFast® SYBR® Green PCR Master Mix (Qiagen). Technical duplicates of this reaction mix were then analyzed on a Corbett Rotor-Gene 6000 real-time PCR cycler. Finally, the copy numbers of the respective tRFs were quantified by standard curves of the individual primer pair amplicons. As normalizers the miRNAs miR25, miR532 and miR99a were used. The boxplots were created with R using the R packages *ggplot2* and *Cairo* (version 3.4.3). qPCR primer sequences are available in Supplementary Table Supplemental_Table_S3.

### RT-PCR quantification of miRanda/piRanha predicted targets

Potential target transcripts of the tRFs were predicted using the algorithm miRanda (version 3.3a) that bases on miRNA targeting rules (Enright et al. 2003) and a self-developed software named piRanha (version 0.0.0) that bases on piRNA targeting rules (Zhang, D. et al. 2018). The reference transcriptome was downloaded from the Ensembl database (release 94). The custom Perl scripts MRscript.pl and PRscript.pl were applied to extract and sort the transcript IDs from the output files by the respective miRanda and piRanha score, which is the sum of binding energies of all alignments with the tRF seed. It is assumed that the lower the miRanda or piRanha score (i.e. free energy of the alignments) of a transcript, the more likely it is a target of the respective tRF. For further analysis, only transcripts that have a TPM value above 0.2 in HEK293 cells were considered. Therefore the RNA sequencing datasets SRR629569 and SRR629570 were downloaded with NCBI’s fastq-dump (-I --split-files --gzip) and uploaded to the Galaxy server (usegalaxy.org), where they were mapped to the human genome (Galaxy hg38) using RNA STAR (Galaxy Tool Version 2.6.0b-1). Transcript wise counting was performed with featureCounts (Galaxy Tool Version 1.6.0.6) on basis of an Ensemble GTF-file (Homo_sapiens.GRCh38.90), which had been converted to UCSC coordinates using the File Chameleon tool of Ensembl. The generated count tables and gene length files were used to calculate the mean TPM values and select the expressed transcripts with R using the R packages *biomaRt* and *stats* (version 3.4.3). For each tRF and algorithm, the five transcripts with the lowest miRanda or piRanha scores were chosen for RT-PCR quantification. Primers with the length of ~20 nt were designed to be either exon-junction spanning or to include intronic regions that are bigger than 700 bp. Furthermore, only primers that exclusively amplify the same amplicon from the different splicing isoforms that are potential targets were taken into account. For each primer pair a test PCR with cDNA from untransfected HEK293T cells was performed to check the amplicon quality and length on an agarose gel. In order to compare the relative copy number of the selected potential tRF targets in tRF-mimic and control transfected cells, the respective total RNA was extracted with TRI Reagent™ (Thermo Fisher), reversely transcribed and quantified as described for the poly-A tailed RNA above. The housekeeping genes *ACTB* and *RPS18* were used as normalizers to calculate the relative expression by means of the delta-delta-C_T_ method. qPCR primer sequences are available in Supplementary Table Supplemental_Table_S3. The boxplots were created with R using the R packages *ggplot2* and *Cairo* (version 3.4.3).

### RNA sequencing, data processing and differential expression analysis

Total RNA isolated from tRF-mimic, tRF-antisense and control transfected cells was send for library construction and paired-end sequencing to BGI (Hong Kong). On average 35 million paired-end reads were obtained. Using the online platform Galaxy (usegalaxy.org) the reads were first mapped to the human genome (Galaxy hg38) by RNA STAR (Galaxy Tool Version 2.6.0b-1). Afterwards, gene wise counting was performed with featureCounts (Galaxy Tool Version 1.6.0.6) on basis of the above mentioned GTF-file. Based on the generated count tables, DESeq2 (Galaxy Tool Version 2.11.40.2) was used to identify differentially expressed genes (adjusted p-value < 0.01). Analysis of the *Drosophila* datasets (Study: PRJNA378597, Run accessions: SRR6930617, SRR6930619, SRR6930621; (Luo et al. 2018)) was likewise conducted using the organism-specific files. Volcano plots based on the DESeq2 result file and cumulative distribution plots based on the DESeq2 result file with thresholds of —80 for miRanda scores and −40 for piRanha scores were created with R using the R packages *ggplot2*, *ggrepel* and *Cairo* (version 3.4.3).

### Identification of targeting patterns using a k-mer mapping approach

tRF sequences with length n were split into all possible k-mers with k=5..n. All k-mers were mapped individually to the 5’ UTR, the CDS and the 3’ UTR of the transcripts of the corresponding organism in reverse complementarity to identify putative target sites. Only the longest transcript per annotated gene was taken into account. The transcriptomes and respective genome annotation files were downloaded from Ensembl database (release 94). Splitting the transcripts into 5’ UTR, CDS and 3’ UTR was performed using the custom Perl scripts select+split_dmel.pl and select+split_hsap.pl. The following numbers of total mismatches (mm, including insertions/deletions) within a k-mer alignment were allowed: For k≥6 1 mm, for k≥12 2 mm, for k≥18 3 mm, for k≥22 4 mm, for k≥26 5 mm. The numbers of allowed insertions/deletions (indel) within a k-mer alignment were: For k≥12 1 indel, for k≥18 2 indels, for k≥22 3 indels. Mismatches were not allowed in the first two or last two position of the alignment. Nested k-mer alignments, i.e. alignments that completely overlapped with larger k-mer alignments, were ignored. k-mer mapping and filtering was performed using the custom Perl script map_kmers.pl.

Gene expression values (fpkm) were calculated based on the featureCounts count tables and genes with an average expression ratio below 1 fpkm were depleted from the DESeq2 result file for further analysis using the R script DESeq2-Analysis-for-get_targets.R. The custom Perl script get_targets_for_DESeq2.pl was then used to count for each tRF and its k-mers defined by start position and length the number of significantly up-/downregulated genes (adjusted p-value < 0.01) that have a corresponding k-mer alignment, and the number of significantly up-/downregulated genes that do not have a corresponding k-mer alignment. The R script get_targets_visualization.R was used to visualize for each transcript region the fraction of targeted as well as not targeted genes that are up-/downregulated. For the individual plots, the script calculates the average values of a 5 nt sliding window for the start position with sliding window increment of 1 nt.

To check whether the transfected tRFs act on a transcriptional or post-transcriptional level, the adapted Perl script select+split4exonjunction_hsap_EJ.pl was used to first split the transcripts into 5’ UTR, CDS and 3’ UTR and then mask all nucleotides, but eight nucleotides at the ultimate exon end (EE), the ultimate exon start (ES) or symmetrically situated around the exon-junction (EJ). Subsequently, 5-mers of the respective tRF sequence was mapped to the masked sequence files using the adapted Perl script map_kmers_EJ.pl. Further analysis and visualization was performed as described above.

### Analysis of potential targets regarding thermodynamically favored alignments with 5’ tRH-Glu-CTC and GO term annotation

Based on the DESeq2 result tables, a Venn diagram of significantly differentially expressed genes (adjusted p-value<0.01) was generated using the R packages *VennDiagram* and *polyclip* (version 3.4.3). Genes that were inversely regulated in the HEK overexpression and the HepG2 inhibition cells were assigned to the groups “potential perish targets” (HEK: log2FC<0; HepG2: log2FC>0) and “potential shelter targets” (HEK: log2FC>0; HepG2: log2FC<0).

Input files for the program RNAup were generated using the custom Perl script RNAup_input.pl, which prints for each potential target gene the sequence of each transcript region (5’ UTR, CDS, 3’ UTR) together with the tRF sequence as fasta-format. RNAup (version 2.4.13) from the ViennaRNA Package (Lorenz et al. 2011) was executed for each input file with the parameters -b -d2 --noLP -c ’S’. Using the custom Perl script merge_RNAupOutput.pl the RNAup output information was merged per transcript region. Using the R script visualize_RNAup.R the respective merged RNAup output files were visualized as a histogram displaying the alignment length, a histogram displaying the alignment start within the tRF and a bar plot displaying the alignment count per position within the tRF.

The list of human protein-coding genes with the gene ontology term “neurogenesis” (GO:0022008) was downloaded from AmiGO 2 (http://amigo.geneontology.org/amigo). Corresponding Ensembl gene identifier for the UniProt identifiers were retrieved from UniProt (https://www.uniprot.org/uploadlists/). To evaluate a potential statistical correlation between the perish or shelter targets of 5’ tRH-Glu-CTC and the GO term “neurogenesis” the chi-squared test was applied.

## Code availability and data deposition

All above mentioned custom Perl and R scripts are freely available at GitHub (github.com/jjehn/tRH-targeting). Sequencing datasets are accessible at NCBI’s sequence read archive (SRA) under the accession numbers SRR10091207 (1^st^ sRNAseq human hippocampus), SRR10091206 (2^nd^ sRNAseq human hippocampus), SRR10091205 (sRNAseq human cortex), SRR10091204 (sRNAseq human cerebellum), SRR10092006 (sRNAseq female chimpanzee hippocampus), SRR10092005 (sRNAseq male chimpanzee hippocampus), SRR10092004 (sRNAseq macaque hippocampus), SRR10082984 (sRNAseq HEK293T cells), and SRR10085693 to SRR10085704 (RNAseq of the HEK293T and HepG2 experiments).

## Acknowledgements

We thank the Wolfrum and Strand lab for kindly providing HEK and HepG2 cells. We further thank René Ketting, Susanne Strand, Daniel Gebert, Christine Barbara Kiefer, and Lena Mazzariello for helpful comments and discussion. Chet C. Sherwood and Cheryl Stimpson from the National Chimpanzee Brain Resource (www.chimpanzeebrain.org, USA) are acknowledged for their great assistance in preparing chimpanzee RNA samples. The chimpanzee samples were supported by the NIH grant NS092988. This work was supported by the International PhD Program (IPP) coordinated by the Institute of Molecular Biology IMB, Mainz, Germany, funded by the Boehringer Ingelheim Foundation.

## Author Contributions

DR and JJ designed the study. VE and CH analyzed the published small RNA sequencing datasets of human tissues. LW and IF generated and analyzed the small RNA libraries of the human hippocampus. RNK dissected the macaque brain and homogenized the hippocampal tissue sample. IF isolated the total RNA from the macaque sample and coordinated the procurement and sequencing of all primate brain samples. JJ analyzed the small RNA sequencing datasets of the cell lines, the primate brain samples and the pig, rat and mouse hippocampus. JJ, SW, and BO performed the laboratory experiments. JJ, JT, and DR analyzed the RNA sequencing data. JJ and DR wrote the manuscript.

