## Supplementary material for "5’ tRNA halves are highly expressed in the primate hippocampus and sequence-specifically regulate gene expression": Figure S1

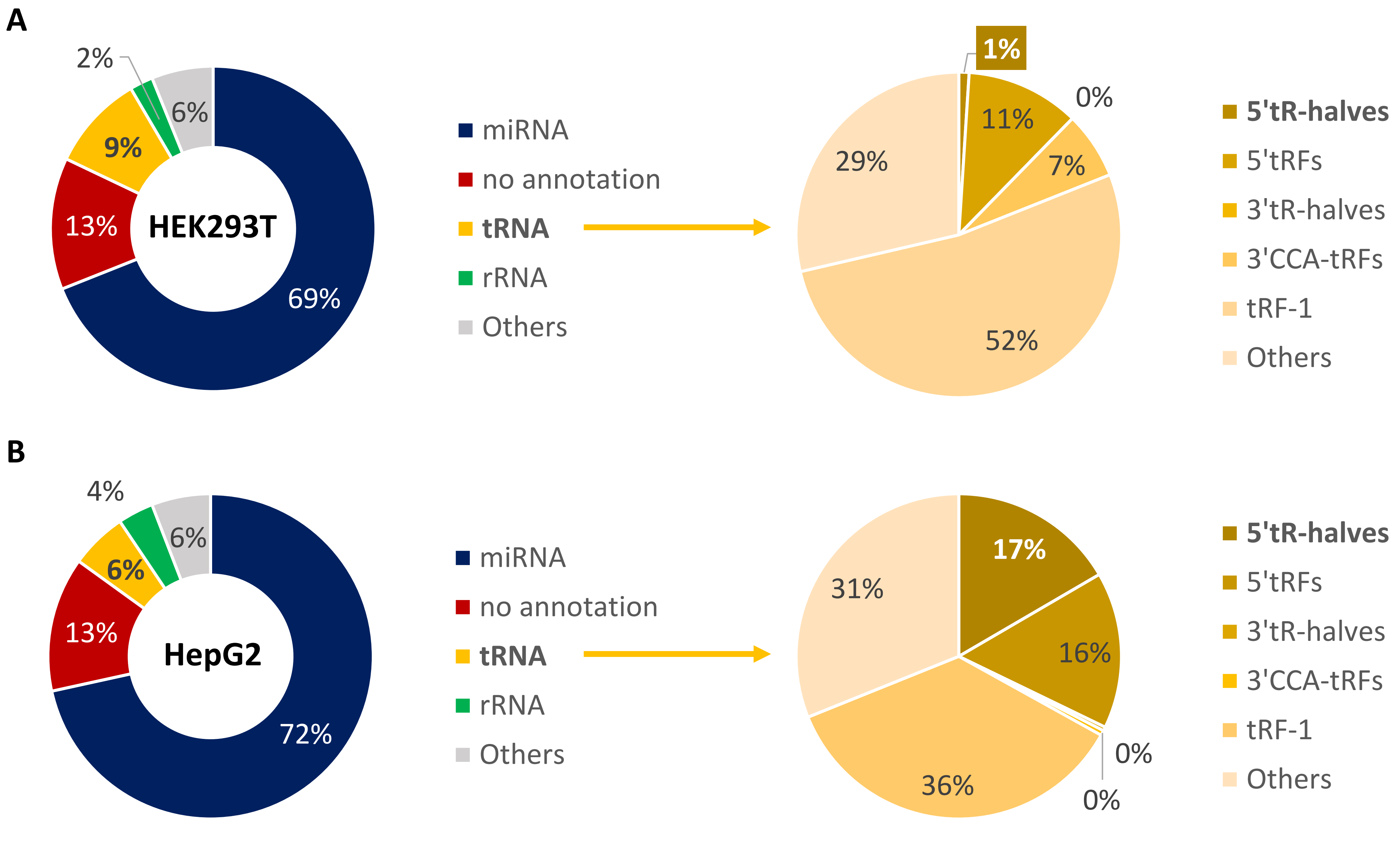


S-Figure 1: Small RNA annotation of mapped reads from small RNA libraries of **(A)** HEK293T cells and **(B)** HepG2 cells.
