## Supplementary material for "5’ tRNA halves are highly expressed in the primate hippocampus and sequence-specifically regulate gene expression": Figure S2

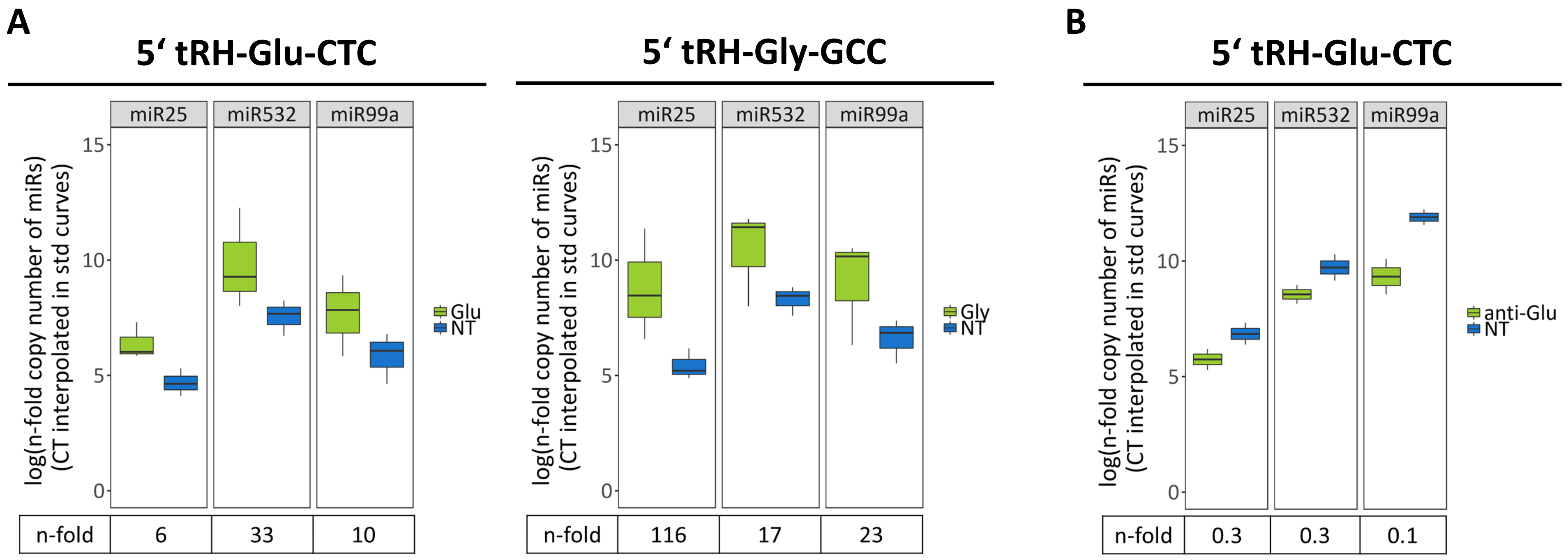


S-Figure 2: **(A)** qPCR quantification of the 5' tRNA-halves Glu-CTC (left) and Gly-GCC (right) in HEK293T cells that were transfected with the synthetic 5’ tRH-mimics (green) or a control non-target siRNA (blue). The three miRNAs miR25, miR532 and miR99a were used as normalizers. The given n-fold change is the ratio between the relative 5’ tRH expression in the overexpression and the control cells. **(B)** qPCR quantification of the 5' tRNA-half Glu-CTC in HepG2 cells that were transfected with an antisense RNA targeting this 5’ tRH (green) or a control non-target siRNA (blue). The three miRNAs miR25, miR532 and miR99a were used as normalizers. The given n fold change is the ratio between the relative 5’ tRH-Glu-CTC expression in the antisense transfected and the control cells.
