## Supplementary material for "5’ tRNA halves are highly expressed in the primate hippocampus and sequence-specifically regulate gene expression": Figure S3

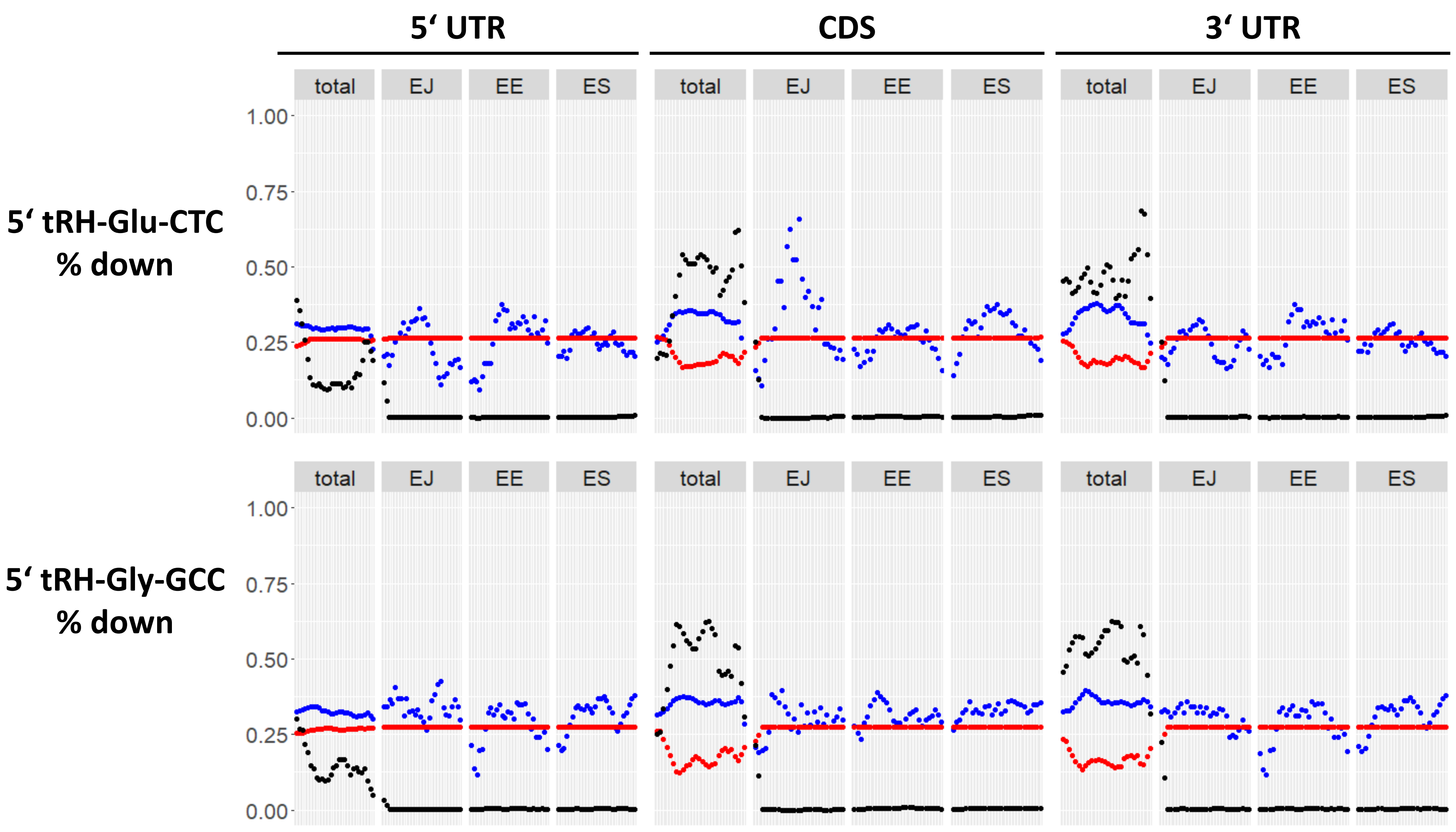


S-Figure 3: Analysis of 5-mer mapping spanning (EJ) or being adjacent to (EE and ES) exon junctions. Displayed is the percentage of transcripts with (blue) or without (red) 5-mer alignment that are downregulated in HEK293T cells upon 5’ tRH mimic transfection.
