## Supplementary material for "5’ tRNA halves are highly expressed in the primate hippocampus and sequence-specifically regulate gene expression": Figure S4

```
human 5' tRh-Glu-CTC     TCCCTGGTGGTCTAGTGGTTAGGATTCGGCGCT
fly   5' tRF-Glu-CTC     T-----ATTGTCTAGTGGTTAGGAT--------
                         *      * ****************
```

S-Figure 4: Sequence alignment between human and fly 5’ tRF Glu-CTC.
