## Supplementary material for "5’ tRNA halves are highly expressed in the primate hippocampus and sequence-specifically regulate gene expression": Table S1

| Tissue | SRA Run Accession# |
|---|---|
| **brain (PFC)** | SRR1635903 |
| **brain (cerebellum)** | SRR1635904 |
| **brain (cerebellum)** | SRR553573 |
| **fibroblast** | SRR4235732 |
| **fibroblast** | SRR4235731 |
| **heart** | SRR4421857 |
| **heart** | SRR4422132 |
| **heart** | SRR553574 |
| **heart** | SRR4422133 |
| **kidney** | SRR1635906 |
| **kidney** | SRR553575 |
| **liver** | SRR1273998 |
| **liver** | SRR1273999 |
| **liver** | SRR1274001 |
| **liver** | SRR4422422 |
| **liver** | SRR4422421 |
| **lung** | SRR1240787 |
| **lung** | SRR1240796 |
| **lung** | SRR1240797 |
| **muscle** | SRR1820680 |
| **muscle** | SRR1820682 |
| **muscle** | SRR1820684 |
| **muscle** | SRR1820686 |
| **muscle** | SRR1820688 |
| **muscle** | SRR1820690 |
| **ovary** | SRR4422259 |
| **ovary** | SRR4422260 |
| **pancreas** | SRR4421754 |
| **pancreas** | SRR4422182 |
| **pancreas** | SRR4422183 |
| **pancreas** | SRR4421755 |
| **prostate** | SRR4421462 |
| **prostate** | SRR4421463 |
| **prostate** | SRR4421707 |
| **prostate** | SRR4421708 |
| **skin** | SRR4421488 |
| **skin** | SRR4421489 |
| **skin** | SRR4422314 |
| **skin** | SRR4422315 |
| **testis** | SRR4422669 |
| **testis** | SRR4422670 |
| **testis** | SRR553576 |
| **testis** | SRR4422476 |
| **testis** | SRR4422477 |
| **thyroid** | SRR4421964 |
| **thyroid** | SRR4421965 |
| **uterus** | SRR4421536 |
| **uterus** | SRR4421537 |
| **uterus** | SRR4421943 |
| **uterus** | SRR4421944 |
