## Supplementary material for "5’ tRNA halves are highly expressed in the primate hippocampus and sequence-specifically regulate gene expression": Table S2

| RNA name | RNA type | Sequence from 5’ to 3’ |
|---|---|---|
| **5’ tRH-Glu-CTC** | RNA | UCCCUGGUGGUCUAGUGGUUAGGAUUCGGCGCU |
| **5’ tRH-Gly-GCC** | RNA | GCAUUGGUGGUUCAGUGGUAGAAUUCUCGCCU |
| **anti-5' tRH-Glu-CTC** | 2’-OMe-RNA | AGCGCCGAAUCCUAACCACUAGACCACCAGGGA |
