## Supplementary material for "5’ tRNA halves are highly expressed in the primate hippocampus and sequence-specifically regulate gene expression": Table S3

| Primer name | Sequence from 5' to 3' | cDNA type |
|---|---|---|
| 5' tRH-Glu-CTC-fwd | TCCCTGGTGGTCTAGTGGTTAG | polyadenylated small RNAs |
| 5' tRH-Gly-GCC-fwd | GCATTGGTGGTTCAGTGGTAG | |
| miR25-fwd | CATTGCACTTGTCTCGGTCTG | |
| miR-532-fwd | CATGCCTTGAGTGTAGGACCGT | |
| miR99a-fwd | AACCCGTAGATCCGATCTTGT | |
| PCR-against-RT-PolyT-rev | CGAATTCTAGAGCTCGAGGCAGG | |
| KIAA1109_fwd | ATCATTTTTCGGTGGTGGAA | Polyadenylated transcripts |
| KIAA1109_rev | CAACTCTTGAAGGCGTCCAT | |
| VPS13D_fwd | CTTACAGGGCAGCATTGGGA | |
| VPS13D_rev | CTGCCTGGAAACGCTGAGTA | |
| SYNE1_fwd | TTTGGAGGCCTGGATAGTGG | |
| SYNE1_rev | AGATCAGAGACCAATGGCGG | |
| FAT1_fwd | CGAGGCATTTGATCCAGATT | |
| FAT1_rev | TCGGTCTAGCTTCCTTGACG | |
| JAG1_fwd | CGATGAATGTGCCAGCAACC | |
| JAG1_rev | CCTTCAGGTGTGTCGTTGGA | |
| RIF1_fwd | TTCTGGAATGCCACTTTTGC | |
| RIF1_rev | CCACTGGATTCCTCCATCAT | |
| HDAC4_fwd | GCAGCACATGGTCTTACTGG | |
| HDAC4_rev | CTGGAACTGCTGCTTGTGTT | |
| IGSF8_fwd | GAAGGTGGCATCCAGAACAT | |
| IGSF8_rev | GGTACACTGTGCCTCCTGCT | |
| HEG1_fwd | AGGAGCGGCTCTTCAAGTAG | |
| HEG1_rev | TGGATGGCAGGTGAAGACTT | |
| RUNX2_fwd | CTGTGGTTACTGTCATGGCG | |
| RUNX2_rev | AGGTAGCTACTTGGGGAGGA | |
| OBSCN_fwd | AGGGCCGAAAATACATCCTG | |
| OBSCN_rev | GACCACATCATACTTCTGGC | |
| HSPG2_fwd | ACTTCATCTCCTTCGGCCTC | |
| HSPG2_rev | TTCCTCGTTCAGATCCAGGC | |
| KLC1_fwd | TGTTCACAAACAGAGGGTGG | |
| KLC1_rev | GCTGCTGTCGTTTTCCACAA | |
| LAMA5_fwd | CAGTACTGTGACATCTGCAC | |
| LAMA5_rev | GGCAAACTTGATGAGGACGT | |
| PLEC_fwd | GAGAAGGTCTTGGCCCTACC | |
| PLEC_rev | ACCAGGCTGATGGTCTTGAG | |
| COL4A3_fwd | TGGCCAGAAAGGATTCACAG | |
| COL4A3_rev | GTACACCGACAAGTCCGTAA | |
| EMILIN3_fwd | CCTCCCGCTACAGTCTCTAC | |
| EMILIN3_rev | CCCATCTACACTGCCGGTAT | |
| TMEM69_fwd | GCCTAGGAACCAATTAGCGC | |
| TMEM69_rev | CTTCTGGATGAAGCGAAGCA | |
| INAVA_fwd | GTGTCCGAGGAGCTCAAGT | |
| INAVA_rev | GATCTCCGCAGCACACAAAC | |
| TMEM8A_fwd | CTGTGCATCCTCAGCTACGA | |
| TMEM8A_rev | TGGAGGCCATGATCACGAAG | |
| ACTB_fwd | CGAGCACAGAGCCTCGCCTTT | |
| ACTB_rev | CATGCCCACCATCACGCCCTGG | |
| RPS18_fwd | GCGGCGGAAAATAGCCTTTG | |
| RPS18_rev | GGATCTTGTACTGGCGTGGA | |
